# Comprehensive analysis of metabolic reprogramming-related gene signaturesmetabolic reprogramming related gene signature for predicting ovarian cancer prognosis, immune landscape, and potential treatment options

**DOI:** 10.1101/2025.08.28.672868

**Authors:** Rendong Han, Enhui Guo, Zhen Li, Huansheng Zhou, Bei Liu, Yan Wang, Xiaomei Hou, FuMin Zheng, Yanan Xu, Jianhong Yu

## Abstract

Ovarian cancer (OV) is the most lethal gynecologic malignancy. Metabolic reprogramming is a distinctive feature of cancer and is associated with tumorigenesis and progression. It could be a potential therapeutic target for cancer treatments and a biomarker for assessing cancer prognosis. In this study, we identified metabolic reprogramming-related differentially expressed genes (MRRDEGs) in OV through differential gene expression analysis and conducted a comprehensive characterization of these MRRDEGs. Based on the MRRDEGs, we constructed an effective prognostic risk model including five model genes for OV. The risk score was a valid independent prognostic factor that could more accurately predict the survival of OV patients. It could classify OV patients into distinct risk groups with significant differences in survival. We observed significant differences between risk groups in biological pathway activity, immune cell infiltration patterns, and immunotherapy responses. Specifically, the low-risk group demonstrated superior immunotherapy response compared to the high-risk group. These findings significantly advance our understanding of the relationship between metabolic reprogramming and OV pathogenesis, progression, prognosis, and immunotherapy response, laying a foundation for developing novel biomarkers and therapeutic targets in the future and providing an important reference for the formulation of precision medicine strategies.

## 1. Introduction

Ovarian cancer (OV) ranks as the third most prevalent gynecological malignancy worldwide, yet it carries the highest mortality rate among all female reproductive system cancers (1). Global cancer statistics reveal an annual incidence of 313,959 OV cases and 207,252 deaths (2), with delayed diagnosis being a major contributor to its poor prognosis. Approximately 66% of OV patients are diagnosed with FIGO (International Federation of Gynaecological Oncology) stage III or IV, and the 5-year survival rates are only 41% and 20%, respectively (3). The standard first-line treatment includes primary cytoreductive surgery (PCS) with platinum-based chemotherapy, supplemented by angiogenesis inhibitors (e.g., bevacizumab) and/or poly (ADP-ribose) polymerase inhibitors (PARPi) as maintenance therapy (1, 4). Despite these interventions, most patients experience disease recurrence (2). The development of resistance not only to chemotherapeutic agents but also to currently approved targeted therapies (bevacizumab and PARPi) highlights the challenges in achieving complete remission in OV (3, 5). Currently, the 5-year survival rate for OV has not improved significantly, and there are still no suitable models or biomarkers to predict the prognosis of OV.

Metabolic reprogramming, a well-recognized hallmark of cancer (including OV), arises as an adaptive response to nutrient deprivation and hypoxia during tumor progression (6, 7). This metabolic rewiring provides essential biomolecules and energy to support malignant proliferation (8). Furthermore, it drives tumor aggressiveness through microenvironmental metabolic competition, contributing to treatment resistance and metastatic dissemination. Recent advances over the past decade have significantly advanced our understanding of cancer metabolism (6), with some metabolic reprogramming activities already being utilized for diagnosis, monitoring, and therapy in oncology (9). In OV specifically, the Warburg effect (aerobic glycolysis) has been demonstrated as critical for tumor initiation and progression (10). Additional metabolic alterations involving amino acid, lipid, and purine metabolism have also been reported (9, 11, 12). Despite these insights, the precise mechanisms governing metabolic reprogramming in OV remain incompletely characterized. Our study aims to systematically characterize metabolic alterations in OV and assess their potential role in ovarian carcinogenesis, as well as their utility for prognostic prediction and therapeutic targeting.

Advances in immunotherapy are further revolutionizing the therapeutic strategies for OV (1). Immune checkpoint inhibitors (ICIs), a breakthrough in cancer treatment, work by reactivating anti-tumor immune responses suppressed by tumor cells (13). While ICIs have shown promising antitumor effects in gynecological cancers, only a small subset of OV patients benefit from immunotherapy. Thus, identifying predictive biomarkers for immunotherapy response is critical to optimize its future application in OV. The limited efficacy is likely attributed to the highly immunosuppressive tumor microenvironment (TME) and its associated alterations (13). Metabolic reprogramming, a key feature of the TME, has been shown to promote immune evasion and hinder immunosurveillance (14, 15). Therefore, it is necessary to study aberrant metabolic reprogramming in OV and the TME to develop biomarkers for predicting immunotherapy response. Furthermore, targeting these metabolic alterations might enhance therapeutic efficacy and overcome resistance to immunotherapy.

Current research should focus on addressing critical gaps in OV clinical management, particularly in elucidating disease pathogenesis, developing novel targeted therapies, and establishing reliable predictive biomarkers. Our study aims to systematically analyze metabolic reprogramming in OV, evaluate its role in tumorigenesis and immune microenvironment regulation, and explore its clinical implications for prognosis prediction and therapeutic targeting. We integrated OV-related data to identify metabolic reprogramming-related differentially expressed genes (MRRDEGs) in OV and analyzed their genomic alteration profiles. Functional enrichment analyses were employed to investigate the association between these MRRDEGs and OV. Furthermore, we constructed a prognostic risk model based on MRRDEGs and validated its predictive performance. Comparative analyses of biological pathway activity and immune microenvironment features were conducted across different risk groups. Finally, we performed differential expression verification and correlation analysis of model genes across multiple independent datasets. The workflow for this study is shown in Figure 1.

**Fig.1.**
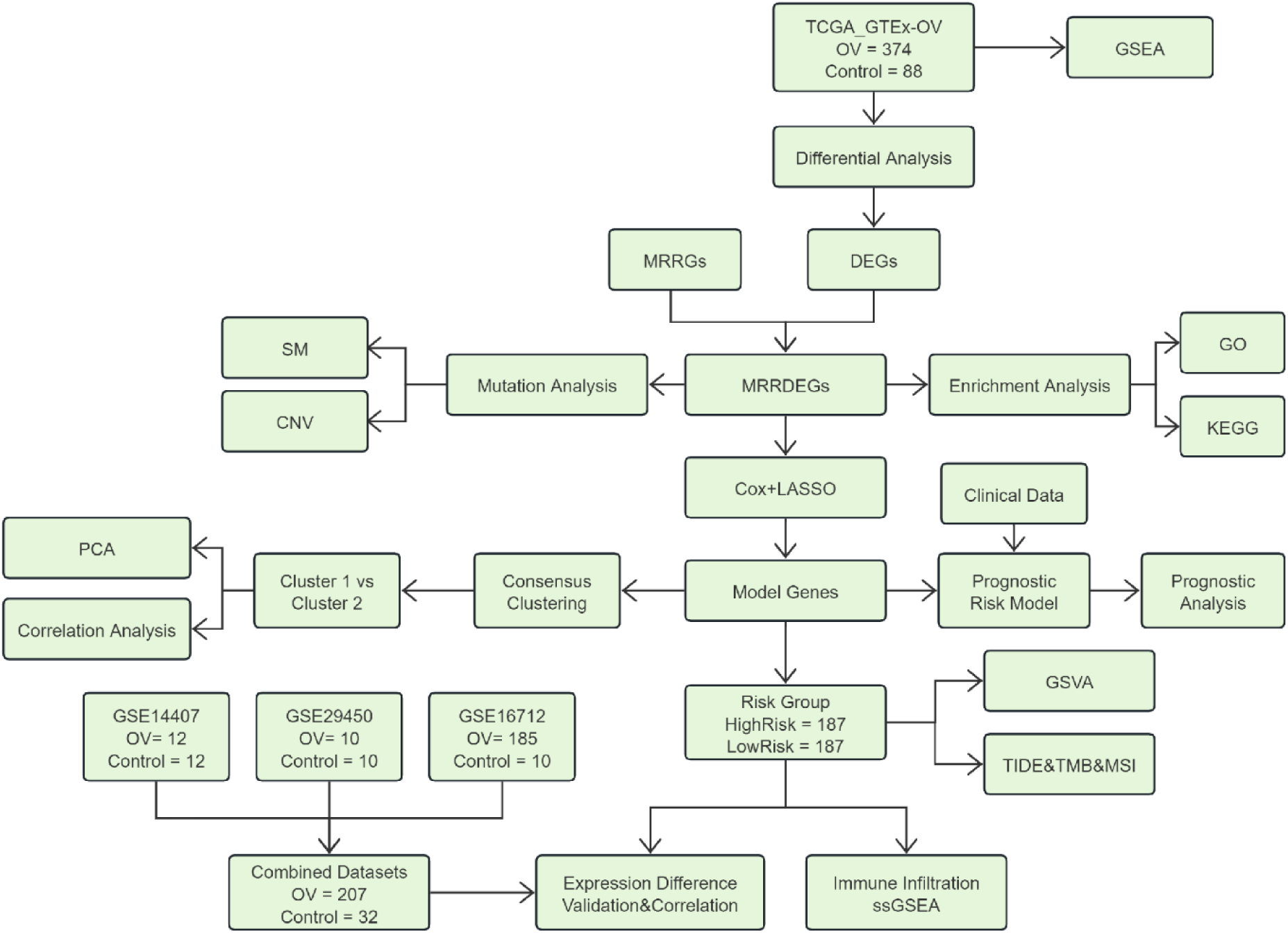
Flow Chart for the Comprehensive Analysis of MRRDEGs. TCGA, The Cancer Genome Atlas; GTEx, Genotype-Tissue Expression; OV, Ovarian cancer; DEGs, Differentially Expressed Genes; MRRGs, Metabolic Reprogramming-Related Genes; MRRDEGs, Metabolic Reprogramming-Related Differentially Expressed Genes; SM, Somatic Mutation; CNV, Copy Number Variation; LASSO, Least Absolute Shrinkage and Selection Operator; GSEA, Gene Set Enrichment Analysis; GO, Gene Ontology; KEGG, Kyoto Encyclopedia of Genes and Genomes; PCA, Principal Component Analysis; GSVA, Gene Set Variation Analysis; TIDE, Tumor Immune Dysfunction and Exclusion; MSI, Microsatellite Instability; TMB, Tumor Mutation Burden; ssGSEA, single-sample Gene Set Enrichment Analysis.

## 2. Materials and Methods

### 2.1 Data Acquisition and Preprocessing

We obtained ovarian cancer (OV) samples from The Cancer Genome Atlas (TCGA) database (https://portal.gdc.cancer.gov/) and control samples from the Genotype-Tissue Expression (GTEx) database (https://www.gtexportal.org/home/) using the R package TCGAbiolinks (16). These datasets were merged to create the TCGA_GTEx-OV dataset, which served as our primary test set. After excluding samples lacking clinical information, we retained 374 OV cases and 88 controls with complete clinical data in counts format. Corresponding clinical metadata were obtained from the UCSC Xena database (17), and the detailed information is shown in Table 1.

**Table 1.** Baseline Table with OV Patients Characteristics.

| Characteristics | Overall |
| --- | --- |
| n | 374 |
| Age, median (IQR) | 59(51, 68) |
| Stage(%) |  |
| Stage I& II | 24 (6.3) |
| Stage III | 293 (78.3) |
| Stage IV | 57 (15.2) |

OV, Ovarian cancer.

Three additional OV datasets (GSE14407 (18), GSE29450 (19), and GSE26712 (20)) were downloaded from the GEO database (https://www.ncbi.nlm.nih.gov/geo/) using the GEOquery package (21) (v2.70.0). All samples were derived from human ovarian tissue, with GSE14407 and GSE29450 using the GPL570 platform and GSE26712 using GPL96. The specific information is shown in Table 2. The GSE14407 dataset contained 12 OV and 12 control samples. The GSE29450 dataset contained 10 OV and 10 control samples. The GSE26712 dataset contained 185 OV and 10 control samples. All OV and control samples were included in this study.

**Table 2.** GEO Microarray Chip Information.

|  | GSE14407 | GSE29450 | GSE26712 |
| --- | --- | --- | --- |
| Platform | GPL570 | GPL570 | GPL96 |
| Species | Homo sapiens | Homo sapiens | Homo sapiens |
| Tissue | Ovarian | Ovarian | Ovarian |
| Samples in OV group | 12 | 10 | 185 |
| Samples in Control group | 12 | 10 | 10 |
| Reference | PMID: 20040092 | PMID:34253236 | PMID: 38057405 |

GEO, Gene Expression Omnibus; OV, Ovarian cancer.

We searched the GeneCards database (22) with “Metabolic Reprogramming” as the search keyword, retained only the protein-coding results, and ultimately obtained 1423 metabolic reprogramming-related genes (MRRGs). Five additional MRRGs were obtained with “Metabolic Reprogramming” as a keyword to search on the PubMed website in the published literature (23). After merging (1,428 total) and removing genes absent from our OV datasets, we retained 1,179 MRRGs for analysis, as presented in Table S1.

The GEO datasets were removed of batch effects using the sva package (24) (v3.50.0), creating a combined dataset of 207 OV and 32 control samples. Data normalization was performed with limma (25) (v3.58.1). Batch effect removal was validated through principal component analysis (PCA) (26).

### 2.2 Differential Expression Analysis

Using limma (25), we compared gene expression between the OV and control groups in TCGA_GTEx-OV, defining differentially expressed genes (DEGs) as those with |logFC| > 4 and adj. p < 0.05. The differential analysis results were visualized as volcano plots using the R package ggplot2 (v3.4.4). MRRGs overlapping with DEGs were designated MRRDEGs, visualized via Venn diagrams. The top 20 MRRDEGs were displayed in heatmaps using pheatmap (v1.0.12).

### 2.3 Genomic Alteration Analysis

To analyze the somatic mutation (SM) in the TCGA_GTEx-OV dataset, the “Masked Somatic Mutation” data was selected and preprocessed using VarScan software. The R package maftools (27) was used to visualize the situation of SM in OV. To analyze the copy number variation (CNV) in the TCGA_GTEx-OV dataset, the “Masked Copy Number Segment” data was selected and analyzed. We visualized the CNV type and frequency using the R package ggplot2 (v3.4.4).

### 2.4 GO and KEGG Pathway Enrichment Analyses

Gene Ontology (GO) enrichment analysis (28) is a commonly used method for conducting large-scale functional enrichment studies, including biological process (BP), cellular component (CC), and molecular function (MF). Kyoto Encyclopedia of Genes and Genomes (KEGG) (29) is a widely used database for storing information related to genomes, biological pathways, diseases, drugs, etc. We performed GO and KEGG pathway enrichment analyses of MRRDEGs using clusterProfiler (v4.10.0) (30). Adj. p < 0.05 and FDR value (q-value) < 0.25 were considered statistically significant thresholds. The p-values were corrected using Benjamini-Hochberg (BH).

### 2.5 Gene Set Enrichment Analysis (GSEA)

GSEA (31) was performed to identify significantly enriched gene sets associated with the phenotype using pre-defined molecular signatures. In this study, the genes from the TCGA_GTEx-OV dataset were ranked by logFC values and analyzed using clusterProfiler with the following parameters: random seed = 2020, minimum gene set size = 10, maximum gene set size = 500. The c2.cp.all.v2022.1.Hs.symbols.gmt gene set from MSigDB (32) was utilized, with significance determined at p < 0.05.

### 2.6 Construction of Prognostic Risk Model

We developed a prognostic risk model for OV using the TCGA_GTEx-OV dataset through a multi-step analytical approach. Least Absolute Shrinkage and Selection Operator (LASSO) regression analysis was performed using the glmnet package (33) (v4.1-8) with family = “cox” specification, incorporating MRRDEGs identified from univariate Cox analysis. LASSO Coefficient Path Plot and Cross-Validation Curve were used to visualize the results of the LASSO regression analysis. Subsequent multivariate Cox regression analysis integrated clinical variables to identify final prognostic model genes, with results presented via a Forest Plot. The risk score was calculated based on the risk coefficient of multivariate Cox regression analysis, and the following formula calculates the risk score:

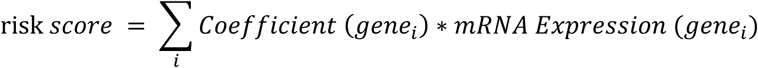

### 2.7 Prognostic Risk Model Validation

We evaluated the prognostic performance of our risk model using multiple approaches. Kaplan-Meier (KM) survival analysis was performed using the survival package (v3.5-7) to compare overall survival (OS) between high- and low-risk groups stratified by the median risk score (34, 35). Time-dependent receiver operating characteristic curve (ROC) (36) was performed based on risk score and OS using survivalROC package (v1.0.3.1) to assess the model’s predictive accuracy at 1-, 3-, and 5-year intervals. AUC values were interpreted as follows: 0.5-0.7 (low accuracy), 0.7-0.9 (moderate accuracy), and > 0.9 (high accuracy).

In addition, we performed comprehensive validation analyses to evaluate the clinical relevance and prognostic power of the risk score. First, univariate and multivariate Cox proportional hazards regression analyses were performed based on risk score, patient age, and clinical stage. Forest plots visualized the results. A prognostic nomogram (37) was constructed using the rms package (v6.7-1) to integrate the multivariate Cox regression results, enabling prediction of 1-, 3-, and 5-year survival probabilities.

The Calibration Curve analysis was used to compare the consistency between the predicted and observed outcomes and assess the accuracy of the model. Decision curve analysis (DCA) implemented with the ggDCA package (v1.1) evaluated the clinical utility of the nomogram for survival prediction.

### 2.8 Gene Set Variation Analysis (GSVA) of Ovarian Cancer Risk Groups

We performed GSVA (38) to investigate pathway activity differences between high- and low-risk OV groups. This non-parametric, unsupervised method transforms gene-level expression matrices into pathway-level enrichment scores. Using the h.all.v7.4.symbols.gmt gene set from MSigDB database, we calculated pathway enrichment scores for all OV samples in the TCGA_GTEx-OV dataset. Samples were stratified into high-risk and low-risk groups based on median risk scores. Pathway enrichment differences were considered statistically significant at p < 0.05.

### 2.9 Validation and Correlation Analysis of Model Genes

To validate the differential expression of model genes between high- and low-risk groups in OV samples, we performed Mann-Whitney U tests on both the TCGA_GTEx-OV dataset and combined datasets. Results were visualized using comparative boxplots showing expression distributions across risk groups.

To explore the co-expression relationships among the model genes, we performed correlation analysis using Spearman’s rank correlation coefficient. The results were visualized through chord diagrams generated with igraph (v1.6.0) and ggraph (v2.1.0) packages. The absolute value of the correlation coefficient (r) below 0.3 was weak or no correlation, 0.3-0.5 was a weak correlation, 0.5-0.8 was a moderate correlation, and above 0.8 was a strong correlation.

### 2.10 Identification of Molecular Subtypes and Correlation Analysis of Model Genes

We performed consensus clustering analysis (39) based on model genes to identify molecular subtypes of OV samples from the TCGA_GTEx-OV. The ConsensusClusterPlus package (40) (v1.62.0) was used. This robust algorithm employs iterative subsampling (50 iterations with 80% sample proportion) to evaluate cluster stability and determine the optimal number of molecular subtypes. Specifically, the number of clusters was set within the range of 2 to 9, with clusterAlg = “km”, and distance metric = “euclidean”. The optimal k was determined by the consensus matrix, consensus cumulative distribution function (CDF) curve, and Delta area plot.

Differential expression of model genes across subtypes was visualized using a heatmap generated by the R package pheatmap (v1.0.12), and subtype separation was validated via a 3D t-SNE plot. Spearman’s rank correlation coefficients were calculated to assess associations between model genes, with significant correlations (p < 0.05) visualized in a heatmap. The top positive and negative correlations were further plotted as scatter diagrams.

### 2.11 Immune Microenvironment Profiling

The tumor immune landscape was characterized using single-sample gene set enrichment analysis (ssGSEA) (41) to quantify the relative abundance of each immune cell population, including activated CD8+ T cell, activated dendritic cell, gamma-delta T cell, natural killer cell, and various human immune cell subtypes such as regulatory T cell (Treg). Enrichment scores calculated from ssGSEA were used to generate an immune infiltration matrix for all samples. Differential immune cell abundance between high- and low-risk groups was visualized using a comparative boxplot created with the ggplot2 package, followed by identification of significantly different immune populations (p < 0.05). Subsequent correlation analyses included examination of immune cell-immune cell interactions through heatmap visualization (pheatmap package) and evaluation of model gene-immune cell relationships using bubble plots (ggplot2 package).

### 2.12 Immunotherapy Biomarker Assessment

The immunotherapy response potential was evaluated through analysis of three key biomarkers. Tumor Immune Dysfunction and Exclusion (TIDE) scores were obtained from the TIDE web portal (http://tide.dfci.harvard.edu) (42, 43) to predict potential tumor treatment response. Microsatellite instability (MSI) status and tumor mutation burden (TMB) data were acquired from the cBioPortal database (https://www.cbioportal.org/) (44). Differences in TIDE scores, MSI status, and TMB between high- and low-risk groups were statistically assessed using Mann-Whitney U tests to identify significant variations in immunotherapy response potential between risk groups.

### 2.13 Statistical Methods

All statistical analyses were performed using R software (v4.3.0). Continuous variables following normal distributions were compared using Student’s t-test, while non-normally distributed variables were analyzed with the Mann-Whitney U test for two-group comparisons or the Kruskal-Wallis test for multiple group comparisons. Associations between variables were examined using Spearman’s rank correlation analysis. All statistical tests were two-tailed, with p-values < 0.05 considered statistically significant.

## 3. Results

### 3.1 Technology Roadmap

### 3.2 Dataset Integration and Batch Effect Correction

We successfully integrated three GEO datasets (GSE14407, GSE26712, GSE29450) using the sva package for batch effect correction, generating a combined dataset. The boxplots (Fig. 2A-B) showed the normalized distribution of expression values after removing batch effects. And the PCA plots (Fig. 2C-D) showed improved clustering of samples by biological origin rather than dataset source. Concordant results from boxplots and PCA confirmed that batch effects were mitigated.

**Fig.2.**
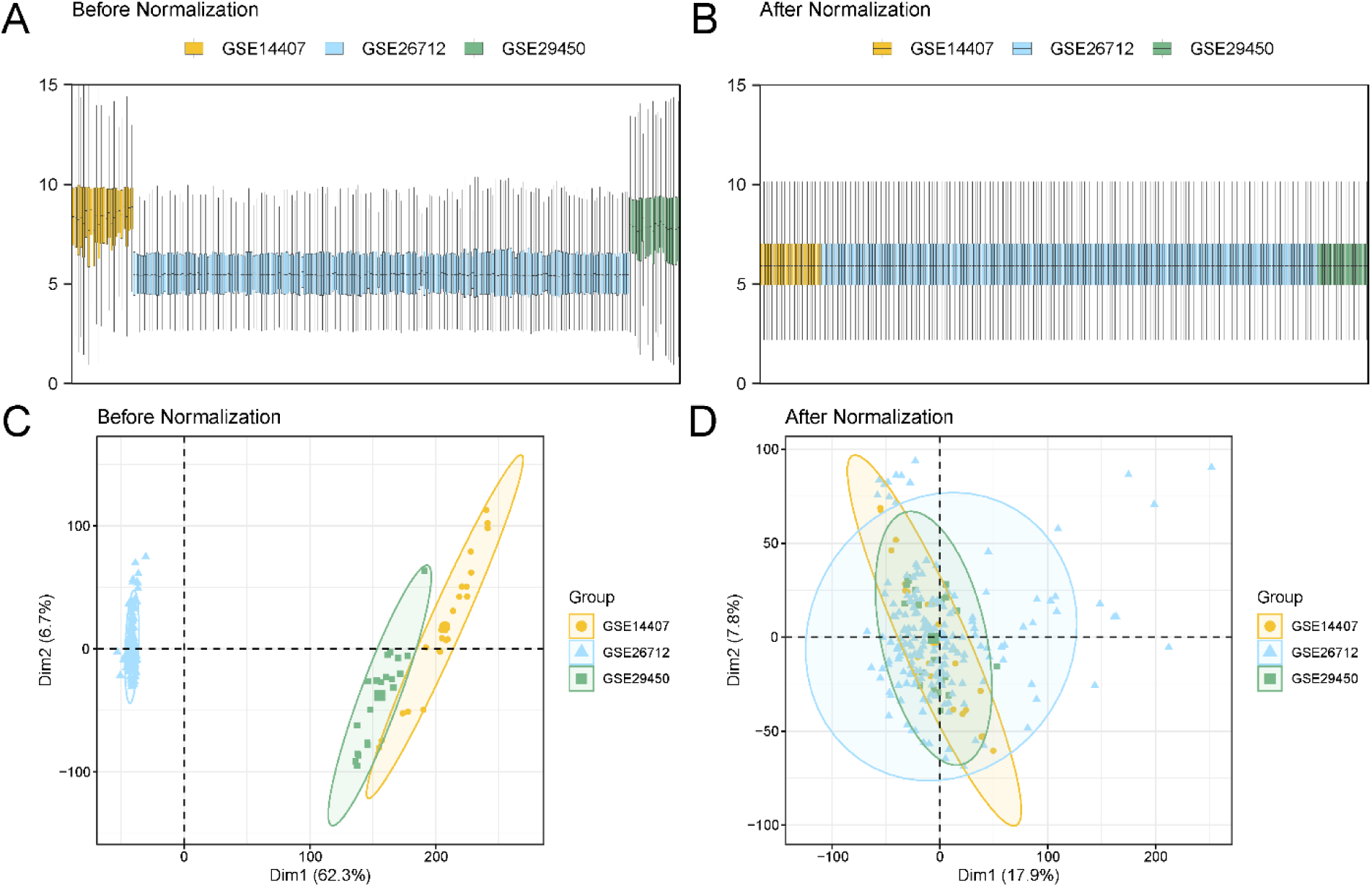
Batch Effects Removal of GSE14407, GSE26712, and GSE29450. A. Expression distribution boxplots before batch correction. B. Expression distribution boxplots after batch correction. C. 2D PCA plot before batch correction. D. 2D PCA plot after batch correction. PCA, Principal Component Analysis. The ovarian cancer dataset GSE14407 is yellow, the ovarian cancer dataset GSE26712 is blue, and the ovarian cancer dataset GSE29450 is green.

### 3.3 Identification of Metabolic Reprogramming-Related Differentially Expressed Genes

Differential expression analysis of the TCGA_GTEx-OV dataset identified 3,107 significant DEGs (|logFC| > 4, adj. p < 0.05), comprising 2,534 upregulated and 573 downregulated genes (Fig. 3A). Intersection with MRRGs yielded 64 high-confidence MRRDEGs (Fig. 3B). The detailed information is shown in Table S2. Heatmap visualization revealed distinct expression patterns of Top20 MRRDEGs between OV and control samples (Fig. 3C).

**Fig.3.**
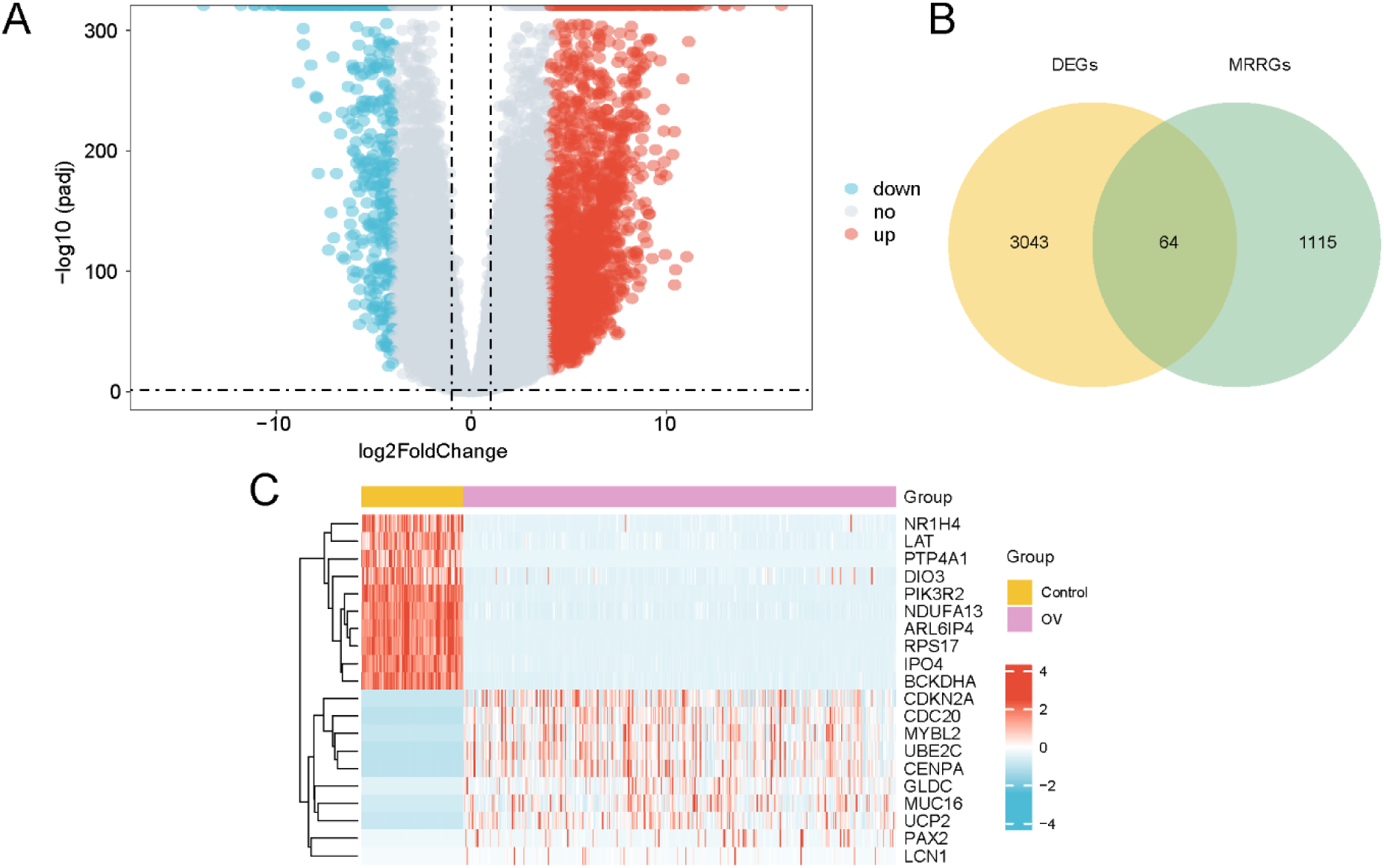
Differential Gene Expression Analysis. A. Volcano plot displaying differentially expressed genes between OV and control groups in the TCGA_GTEx-OV dataset (|logFC| > 4, adj. p < 0.05). B. Venn diagram illustrating the intersection between DEGs and MRRGs. C. Heatmap of MRRDEGs across samples. OV, Ovarian cancer; TCGA, The Cancer Genome Atlas; GTEx, Genotype-Tissue Expression; DEGs, Differentially Expressed Genes; MRRGs, Metabolic Reprogramming-Related Genes; MRRDEGs, Metabolic Reprogramming-Related Differentially Expressed Genes. Pink is the OV group, and yellow is the control group. In the heatmap, red represents high expression, blue represents low expression, and the depth of color represents the degree of expression.

### 3.4 Genomic Alterations in Metabolic Reprogramming-Related Differentially Expressed Genes

SM analysis of all genes revealed nine predominant mutation types in OV, with missense mutations being most frequent (Fig. 4A). Single nucleotide polymorphisms (SNPs) constituted the majority of genetic alterations, with C>T transitions representing the most frequent single nucleotide variant (SNV) in OV. Among MRRDEGs, *MUC16* showed the highest mutation frequency (8%) (Fig. 4B). CNV analysis identified 52 MRRDEGs with significant CNV alterations. The top 20 genes with CNV were visualized through lollipop plots (mutation types, Fig. 4C) and frequency histograms (Fig. 4D).

**Fig.4.**
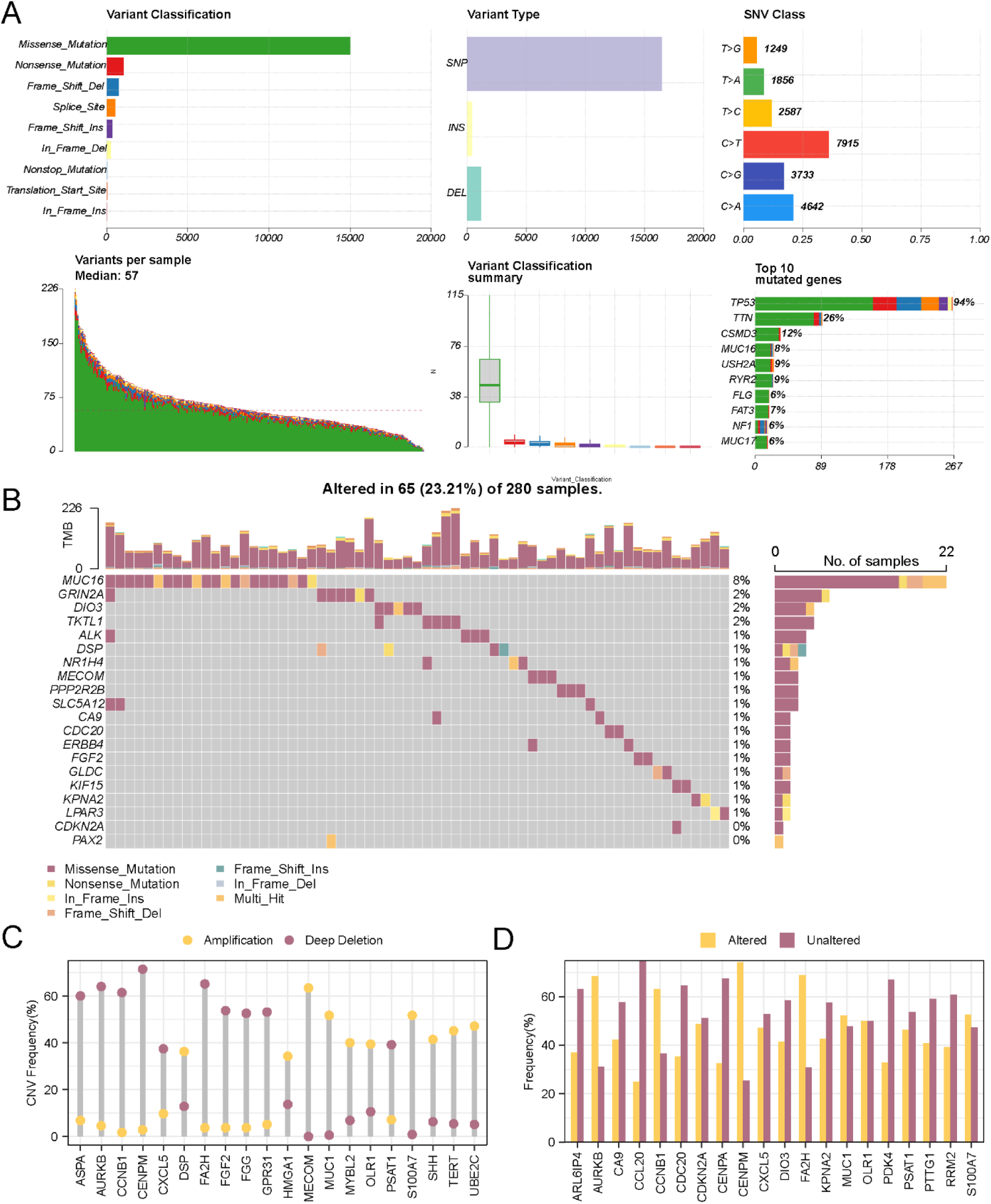
Genomic Alteration Profiles. A. SM landscape in OV. B. Mutation frequency of the top 20 MRRDEGs. C-D. CNV of MRRDEGs in OV: the mutation type lollipop figure (C), and mutation frequency histogram (D). SM, Somatic Mutation; OV, Ovarian cancer; MRRDEGs, Metabolic Reprogramming-Related Differentially Expressed Genes; CNV, Copy Number Variation; SNV, Single Nucleotide Variant; SNP, Single Nucleotide Polymorphism.

### 3.5 GO and KEGG Pathway Enrichment Analyses

GO and KEGG pathway enrichment analyses were performed to elucidate the biological significance of the 64 MRRDEGs in OV (Table 3). The MRRDEGs were significantly enriched in the mitotic nuclear division, regulation of mitotic cell cycle phase transition, regulation of mitotic cell cycle phase transition, regulation of cell cycle phase transition, mitotic cell cycle phase transition, nuclear division, and other BP; chromosomal region, spindle, mitochondrial matrix, kinetochore, condensed chromosome, centromeric region, and other CC. It was also enriched in the Cell Cycle, Cellular senescence, Human T-cell leukemia virus 1 infection, p53 signaling pathway, Gastric cancer, and other biological pathways. The results of GO and KEGG pathway enrichment analyses were visualized through bar graphs (Fig. 5A) and network diagrams (Figs. 5B-D) that illustrate the relationships between molecular entities and functional annotations.

**Fig.5.**
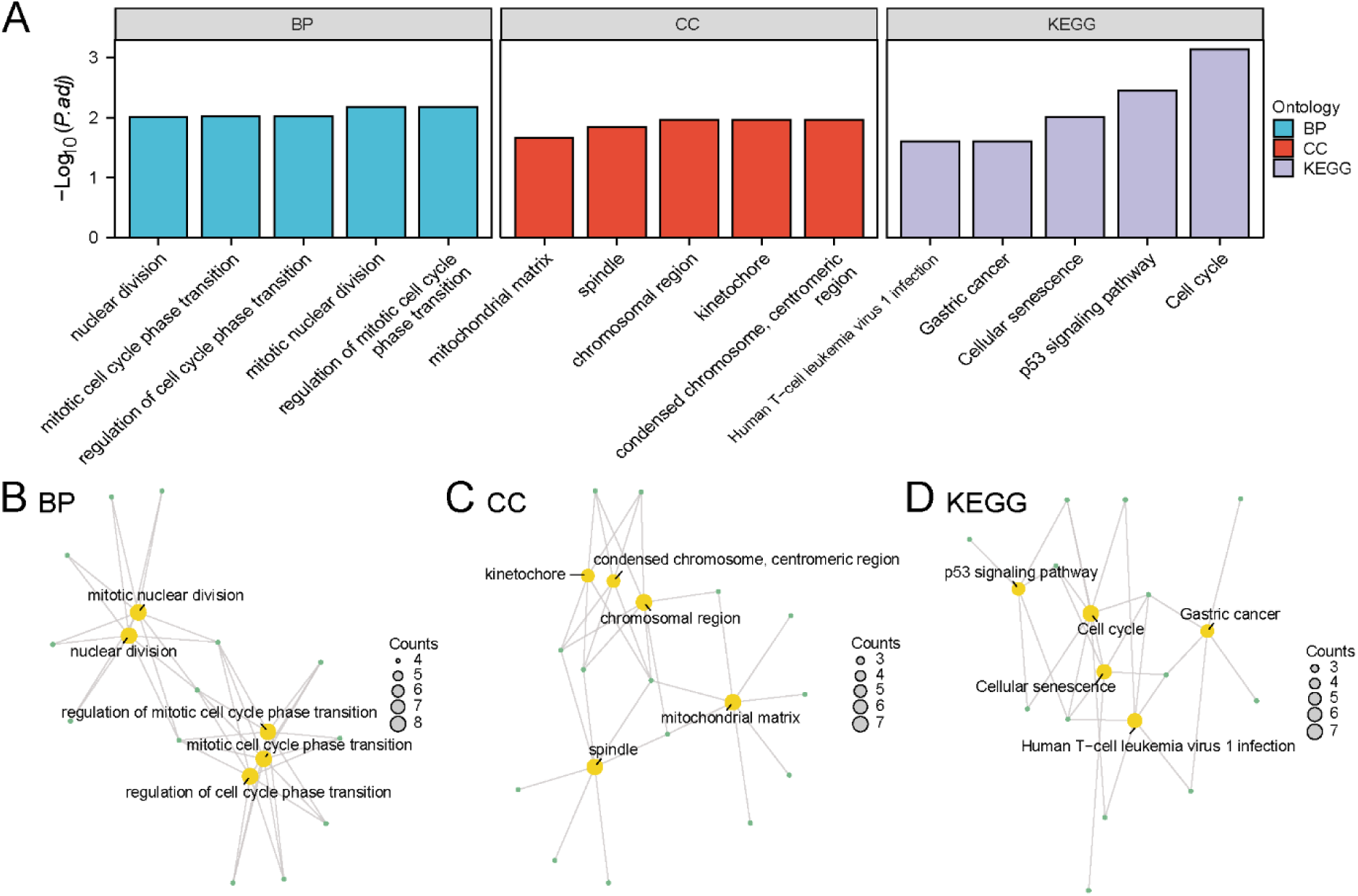
GO and KEGG Pathway Enrichment Analyses of MRRDEGs. A. Bar graph showing significantly enriched GO terms and KEGG pathways. Network diagram of GO and KEGG pathway enrichment analyses results of MRRDEGs: BP (B), CC (C), and KEGG pathways (D). The node size corresponds to the number of associated molecules, with yellow nodes indicating functional terms and green nodes representing genes. The lines represent the relationship between terms and genes. MRRDEGs, Metabolic Reprogramming-Related Differentially Expressed Genes; GO, Gene Ontology; BP, Biological Process; CC, Cellular Component; KEGG, Kyoto Encyclopedia of Genes and Genomes. The screening criteria for GO and KEGG pathway enrichment analyses were adj. p < 0.05 and FDR value (q-value) < 0.25, and the p-value correction method was Benjamini-Hochberg (BH).

**Table 3.** Result of GO and KEGG Pathway Enrichment Analyses for MRRDEGs.

| ONTOLOGY | ID | Description | GeneRatio | BgRatio | p-value | p.adjust | q-value |
| --- | --- | --- | --- | --- | --- | --- | --- |
| BP | GO:0140014 | mitotic nuclear division | 8/63 | 293/18800 | 5.83 e-06 | 6.72 e-03 | 5.08 e-03 |
| BP | GO:1901990 | regulation of mitotic cell cycle phase transition | 8/63 | 321/18800 | 1.13 e-05 | 6.72 e-03 | 5.08 e-03 |
| BP | GO:1901987 | regulation of cell cycle phase transition | 8/63 | 415/18800 | 7.06 e-05 | 9.49 e-03 | 7.17 e-03 |
| BP | GO:0044772 | mitotic cell cycle phase transition | 8/63 | 440/18800 | 1.06 e-04 | 9.66 e-03 | 7.31 e-03 |
| BP | GO:0000280 | nuclear division | 8/63 | 446/18800 | 1.16 e-04 | 9.73 e-03 | 7.35 e-03 |
| CC | GO:0098687 | chromosomal region | 7/64 | 366/19594 | 1.86 e-04 | 1.10 e-02 | 8.57 e-03 |
| CC | GO:0005819 | spindle | 7/64 | 402/19594 | 3.30 e-04 | 1.46 e-02 | 1.14 e-02 |
| CC | GO:0005759 | mitochondrial matrix | 7/64 | 473/19594 | 8.66 e-04 | 2.21 e-02 | 1.72 e-02 |
| CC | GO:0000776 | kinetochore | 5/64 | 146/19594 | 1.15 e-04 | 1.10 e-02 | 8.57 e-03 |
| CC | GO:0000779 | condensed chromosome, centromeric region | 5/64 | 156/19594 | 1.57 e-04 | 1.10 e-02 | 8.57 e-03 |
| KEGG | hsa04110 | Cell cycle | 7/44 | 126/8164 | 4.21 e-06 | 7.32 e-04 | 6.38 e-04 |
| KEGG | hsa04218 | Cellular senescence | 6/44 | 156/8164 | 1.71 e-04 | 9.93 e-03 | 8.65 e-03 |
| KEGG | hsa05166 | Human T-cell leukemia virus 1 infection | 6/44 | 222/8164 | 1.12 e-03 | 2.52 e-02 | 2.20 e-02 |
| KEGG | hsa04115 | p53 signaling pathway | 5/44 | 73/8164 | 4.12 e-05 | 3.58 e-03 | 3.12 e-03 |
| KEGG | hsa05226 | Gastric cancer | 5/44 | 149/8164 | 1.16 e-03 | 2.52 e-02 | 2.20 e-02 |

GO, Gene Ontology; BP, Biological Process; CC, Cellular Component; KEGG, Kyoto Encyclopedia of Genes and Genomes; MRRDEGs, Metabolic Reprogramming-Related Differentially Expressed Genes.

### 3.6 Gene Set Enrichment Analysis (GSEA) in Ovarian Cancer

To investigate the functional implications of global gene expression patterns in OV, we performed GSEA on the TCGA_GTEx-OV dataset. This analysis elucidated the functional relationships between global gene expression patterns and their associated BP, CC, and MF (Fig. 6A). The detailed results are presented in Table 4. The results demonstrated significant enrichment of all genes from the TCGA_GTEx-OV dataset in biological pathways, including: Fceri Mediated Mapk Activation (Fig. 6B), FCGR3A Mediated IL10 Synthesis (Fig. 6C), Fceri Mediated Nf Kb Activation (Fig. 6D), and Ethanol Effects On Histone Modifications (Fig. 6E).

**Fig.6.**
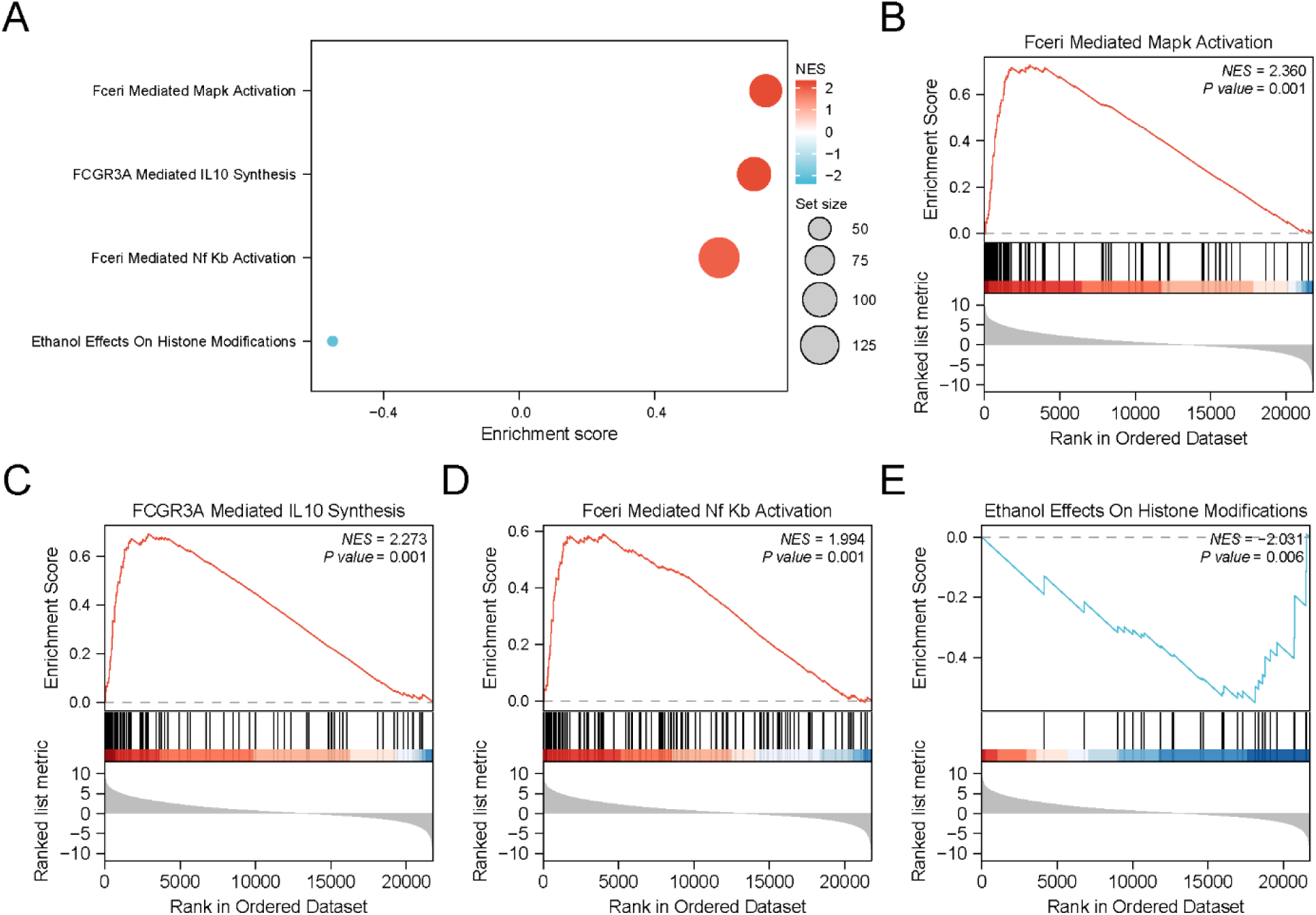
GSEA of TCGA_GTEx-OV Dataset. A. Bubble plot displaying four significantly enriched biological functions identified by GSEA in OV (TCGA_GTEx-OV). B-E. GSEA plots showing significant enrichment in: Fceri Mediated Mapk Activation (B), FCGR3A Mediated IL10 Synthesis (C), Fceri Mediated Nf Kb Activation (D), and Ethanol Effects On Histone Modifications (E). GSEA, Gene Set Enrichment Analysis; TCGA, The Cancer Genome Atlas; GTEx, Genotype-Tissue Expression; OV, Ovarian cancer. The screening criterion of GSEA was p < 0.05.

**Table 4.**
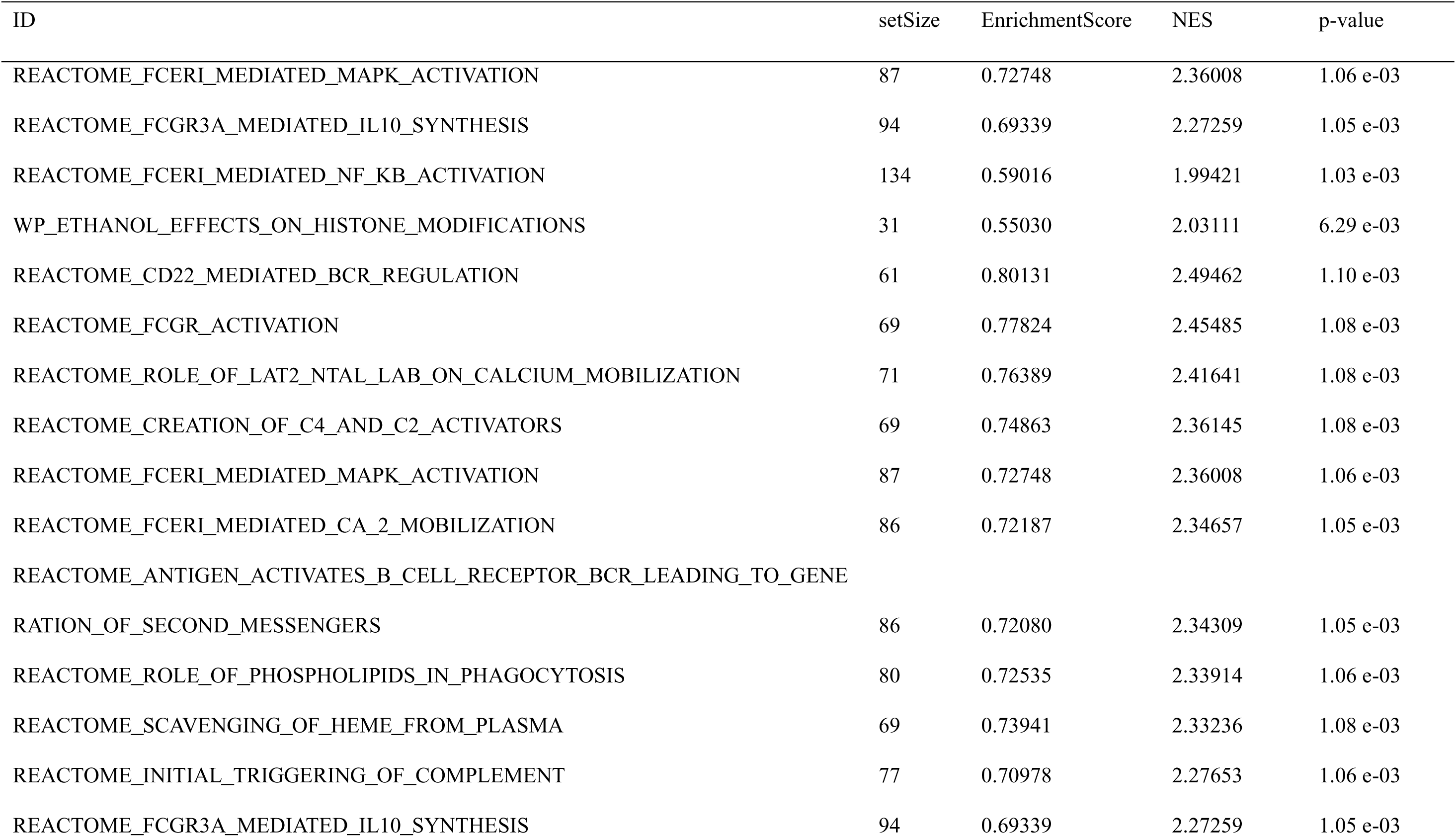
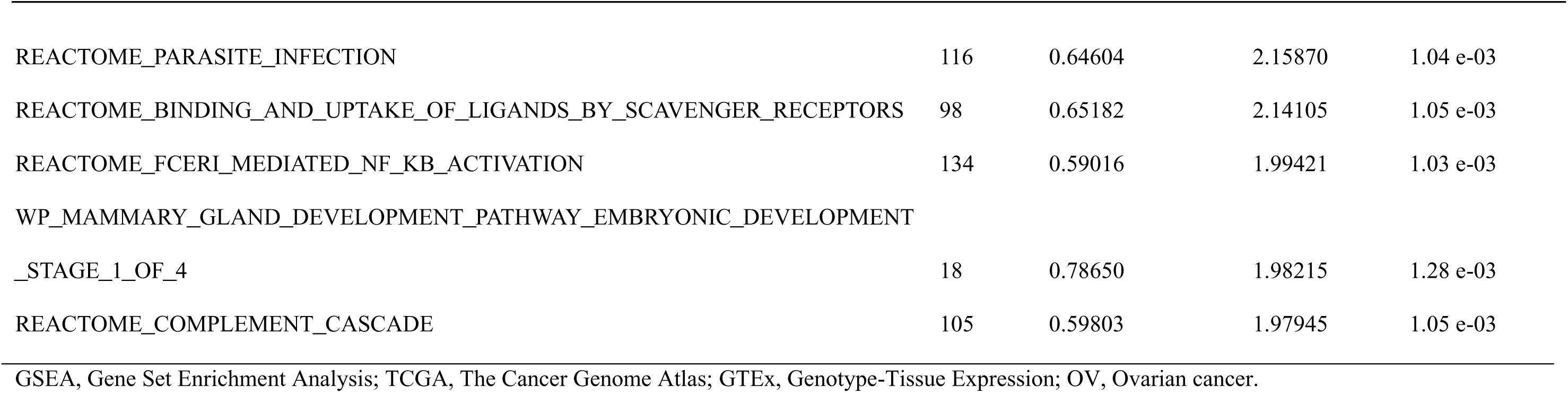
Results of GSEA for TCGA_GTEx-OV.

### 3.7 Construction of Ovarian Cancer Prognostic Risk Model

Our prognostic risk model construction began with univariate Cox regression analysis of 64 MRRDEGs. Five MRRDEGs showed statistical significance (p < 0.10, Fig. 7A). Subsequent LASSO regression analysis (Fig. 7B-C) identified five key prognostic genes, namely Pituitary tumor-transforming gene 1 (*PTTG1*), Lysophosphatidic acid receptor 3 (*LPAR3*), Paired Box Gene 2 (*PAX2*), Pyruvate dehydrogenase kinase 4 (*PDK4*), and Iodothyronine Deiodinase 3 (*DIO3*). Finally, multivariate Cox regression analysis incorporating these five prognostic genes was performed and visualized via a forest plot (Fig. 7D), demonstrating the correlation between the risk score levels and the clinical prognosis. The risk score was calculated as follows:

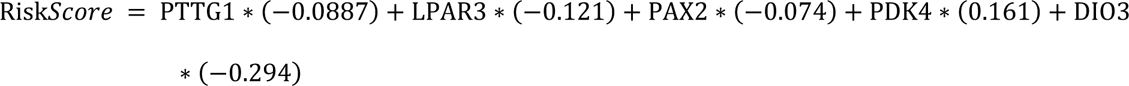

**Fig.7.**
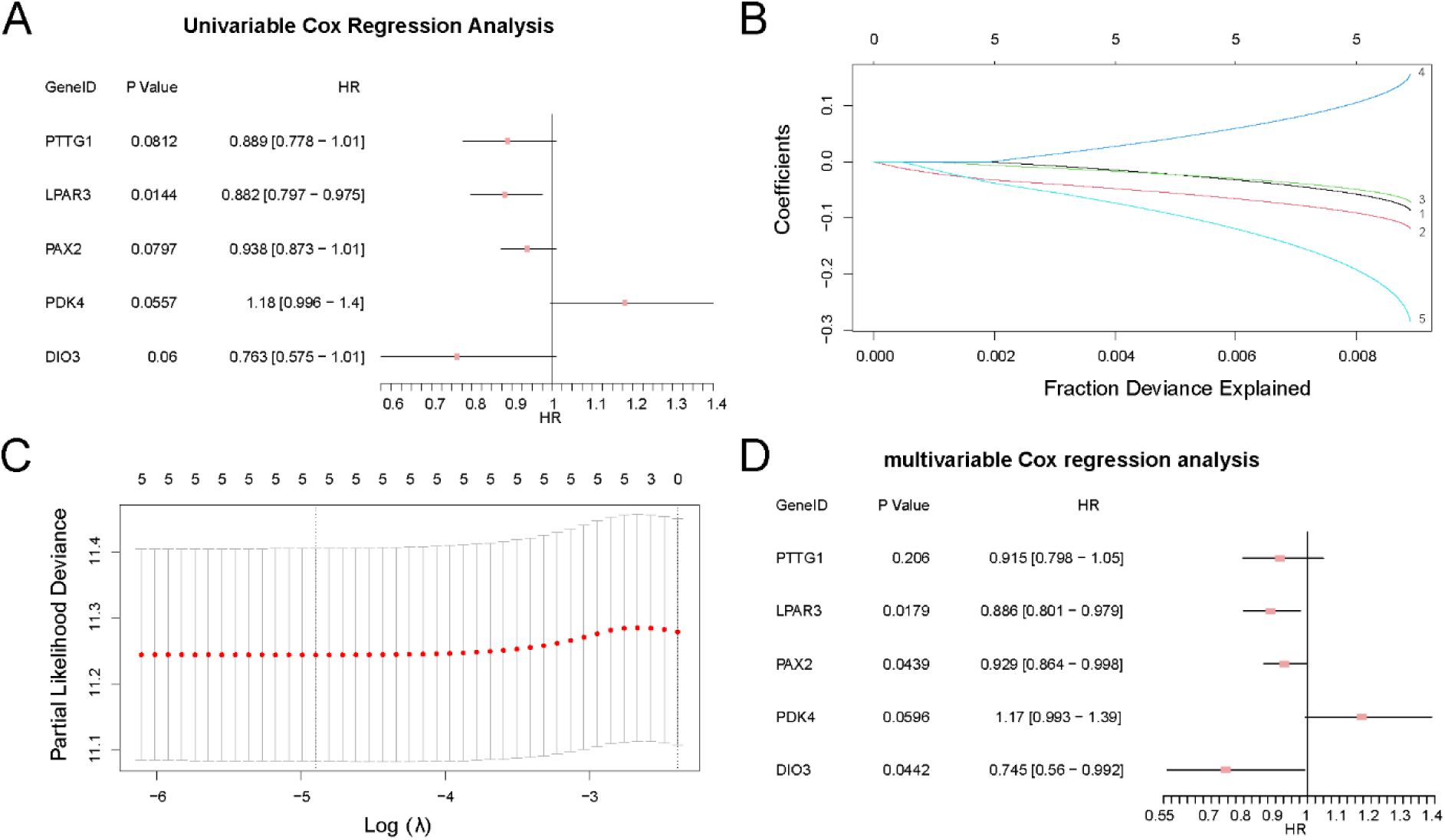
Cox and LASSO Regression Analyses. A. Forest plot of five significant MRRDEGs from univariate Cox regression analysis. B. LASSO coefficient path plot. C. LASSO cross-validation curve. D. Forest plot of multivariate Cox regression analysis. LASSO, Least Absolute Shrinkage and Selection Operator; MRRDEGs, Metabolic Reprogramming-Related Differentially Expressed Genes.

### 3.8 Validation of Ovarian Cancer Prognostic Risk Model

Time-dependent ROC analysis demonstrated that the OV prognostic risk model exhibited low predictive accuracy for 1-, 3-, and 5-year survival (AUC: 0.5-0.7, Fig. 8A). KM analysis revealed highly statistically significant differences in OS between high- and low-risk groups (p < 0.001, Fig. 8B).

**Fig.8.**
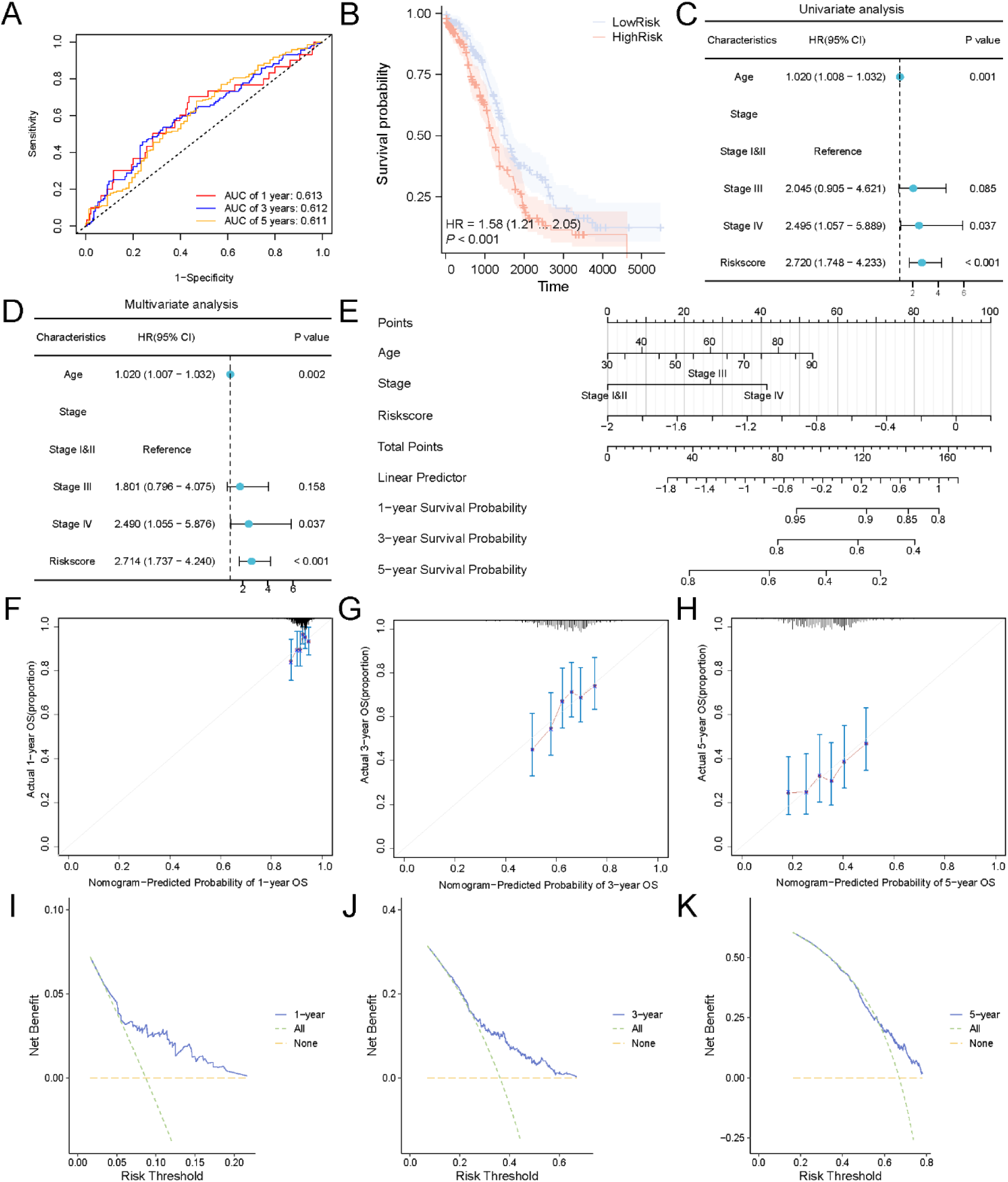
Validation of Prognostic Risk Model. A. Time-dependent ROC curves. B. KM survival analysis between high and low-risk score groups. C. Forest plot in the univariate Cox regression analysis. D. Forest plot in the multivariate Cox regression analysis. E. Prognostic nomogram. F-H. 1-year (F), 3-year (G), and 5-year (H) calibration curves. I-K. 1-year (I), 3-year (J), and 5-year (K) DCA plots. ROC, Receiver Operating Characteristic Curve; AUC, Area Under the Curve; KM, Kaplan-Meier; DCA, Decision Curve Analysis. When AUC > 0.5, it indicates that the expression of the molecule is a trend to promote the occurrence of the event. And the closer the AUC is to 1, the better the diagnostic effect. AUC had lower accuracy in the range of 0.5 to 0.7. A p-value of less than 0.001 was highly statistically significant.

The univariate analysis initially incorporated risk score, age, and clinical stage variables, with all factors showing p < 0.10 subsequently included in multivariate analysis (Fig. 8C-D, Table 5). Both univariate and multivariate Cox analyses identified risk score, age, and clinical stage as significant prognostic factors. The nomogram (Fig. 8E) indicated that risk score contributed substantially more to prognostic prediction than other variables, while clinical stage showed relatively weaker predictive value.

**Table 5.** Results of Univariable and Multivariable Cox Analysis.

| Characteristics | Total(N) | Univariate analysis |  | Multivariate analysis |  |
| --- | --- | --- | --- | --- | --- |
|  |  | Hazard ratio (95% CI) | p-value | Hazard ratio (95% CI) | p-value |
| Age | 374 | 1.020 (1.008-1.032) | 0.001 | 1.020 (1.007-1.032) | 0.002 |
| Stage | 374 |  |  |  |  |
| Stage I&II | 24 | Reference |  | Reference |  |
| Stage III | 293 | 2.045 (0.905-4.621) | 0.085 | 1.801 (0.796-4.075) | 0.158 |
| Stage IV | 57 | 2.495 (1.057-5.889) | 0.037 | 2.490 (1.055-5.876) | 0.037 |
| Riskscore | 374 | 2.720 (1.748-4.233) | < 0.001 | 2.714 (1.737-4.240) | < 0.001 |

The Calibration curves (Fig. 8F-H) showed optimal predictive performance at 1-year follow-up. Finally, DCA (Fig. 8I-K) revealed the greatest clinical utility at 3-year follow-up (3-year > 1-year > 5-year).

### 3.9 Gene Set Variation Analysis (GSVA) of Ovarian Cancer Risk Groups

We performed GSVA to systematically characterize pathway-level differences between high- and low-risk OV groups in the TCGA_GTEx-OV dataset (Table 6). Eighteen pathways showed significant differential enrichment (p < 0.05) between risk groups. The distinct pathway activation patterns between high and low-risk groups were analyzed and visualized by heatmap (Fig. 9A). Mann-Whitney U tests confirmed that these pathway differences between high and low-risk groups were statistically significant (p < 0.05, Fig. 9B).

**Fig.9.**
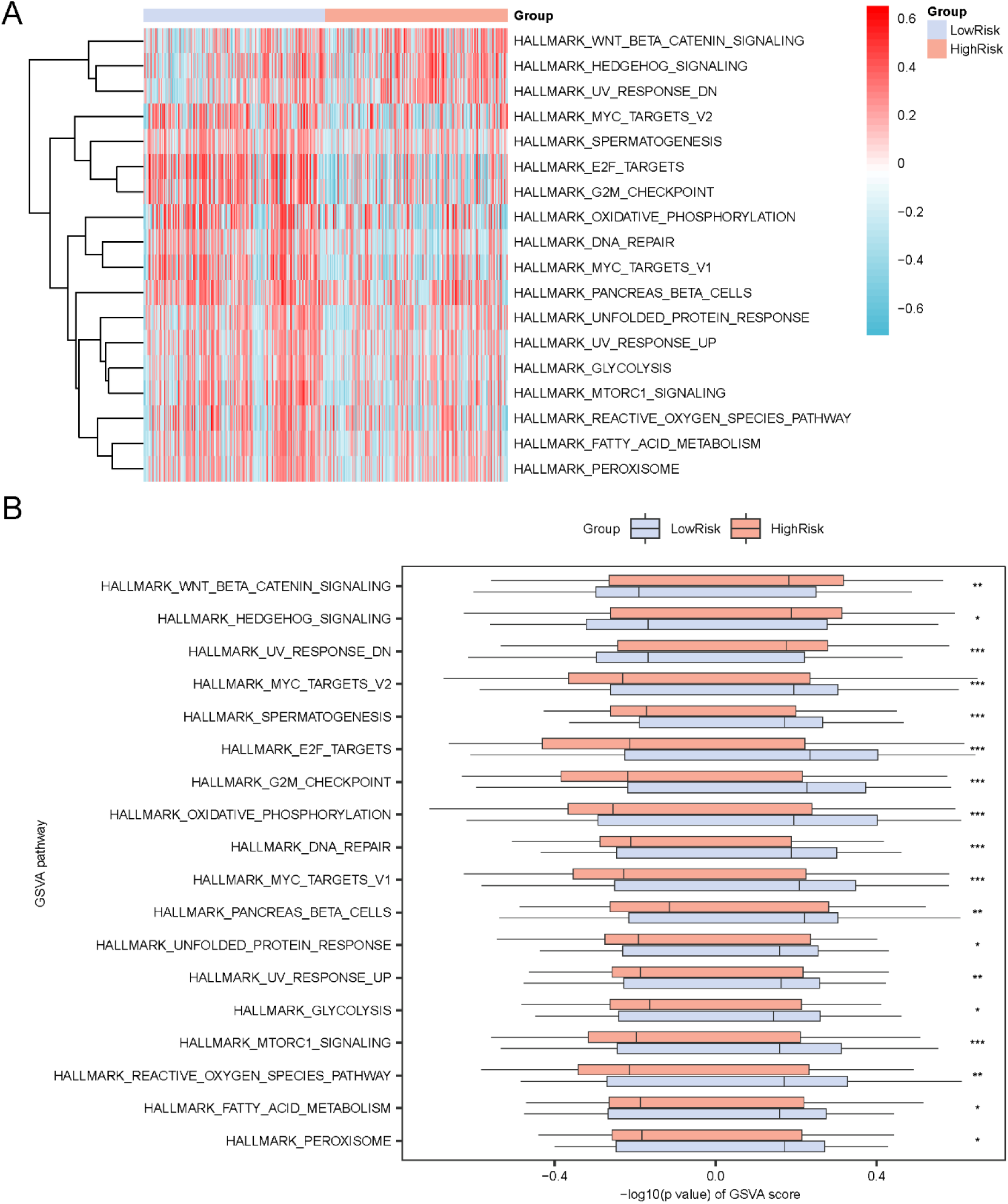
GSVA of OV Risk Groups. A. Heatmap of GSVA scores for 18 significantly enriched pathways. B. Box plots comparing pathway activity between high and low-risk groups. GSVA, Gene Set Variation Analysis; OV, Ovarian cancer. * represents p < 0.05, statistically significant; ** represents p < 0.01, highly statistically significant; *** represents p < 0.001 and highly statistically significant. Light red represents the high-risk group; Light blue represents the low-risk group. Red indicates high enrichment and blue indicates low enrichment in the heatmap. The screening criterion for GSVA was p < 0.05.

**Table 6.** Results of GSVA for OV Risk Groups.

| ID | logFC | AveExpr | p-value | adj. p value |
| --- | --- | --- | --- | --- |
| HALLMARK_E2F_TARGETS | 0.243572 | 0.007777 | 7.62 e-11 | 3.81 e-09 |
| HALLMARK_G2M_CHECKPOINT | 0.205538 | 0.011590 | 5.40 e-09 | 1.35 e-07 |
| HALLMARK_MYC_TARGETS_V1 | 0.181948 | 0.026460 | 6.44 e-08 | 1.07 e-06 |
| HALLMARK_DNA_REPAIR | 0.148199 | 0.037921 | 2.82 e-07 | 3.52 e-06 |
| HALLMARK_OXIDATIVE_PHOSPHORYLATION | 0.177216 | 0.028170 | 1.66 e-06 | 1.66 e-05 |
| HALLMARK_SPERMATOGENESIS | 0.106139 | 0.004731 | 5.32 e-05 | 3.67 e-04 |
| HALLMARK_MYC_TARGETS_V2 | 0.139847 | 0.024325 | 5.35 e-05 | 3.67 e-04 |
| HALLMARK_MTORC1_SIGNALING | 0.124414 | 0.022515 | 5.87 e-05 | 3.67 e-04 |
| HALLMARK_UV_RESPONSE_UP | 0.088434 | 0.016115 | 1.07 e-03 | 5.96 e-03 |
| HALLMARK_UV_RESPONSE_DN | 0.096711 | 0.009415 | 1.32 e-03 | 6.61 e-03 |
| HALLMARK_PANCREAS_BETA_CELLS | 0.091965 | 0.049601 | 2.69 e-03 | 1.22 e-02 |
| HALLMARK_WNT_BETA_CATENIN_SIGNALING | 0.092294 | 0.013688 | 3.55 e-03 | 1.44 e-02 |
| HALLMARK_REACTIVE_OXYGEN_SPECIES_PATHWAY | 0.094470 | 0.028429 | 3.73 e-03 | 1.44 e-02 |
| HALLMARK_PEROXISOME | 0.072405 | 0.016752 | 8.00 e-03 | 2.86 e-02 |
| HALLMARK_UNFOLDED_PROTEIN_RESPONSE | 0.070541 | 0.012584 | 1.32 e-02 | 4.40 e-02 |
| HALLMARK_GLYCOLYSIS | 0.067245 | 0.017721 | 1.47 e-02 | 4.61 e-02 |
| HALLMARK_FATTY_ACID_METABOLISM | 0.066724 | 0.013991 | 1.94 e-02 | 5.71 e-02 |
| HALLMARK_HEDGEHOG_SIGNALING | 0.071583 | 0.002942 | 3.15 e-02 | 8.75 e-02 |

GSVA, Gene Set Variation Analysis; OV, Ovarian cancer.

### 3.10 Validation and Correlation Analysis of Model Genes

To validate the risk-stratification potential of our prognostic model gene signature, we compared the expression patterns of five candidate genes (*PTTG1, LPAR3, PAX2, PDK4, DIO3*) between high- and low-risk OV groups across two independent datasets. All five genes showed highly statistically significant differential expression between the high and low-risk groups in the TCGA_GTEx-OV dataset (p < 0.001, Fig. 10A). Four genes (*PTTG1*, *LPAR3*, *PDK4*, *DIO3*) maintained highly statistically significant differential expression between the high and low-risk groups in the combined dataset (p < 0.001, Fig. 10B). *PAX2* expression showed no significant intergroup difference (p > 0.01).

**Fig.10.**
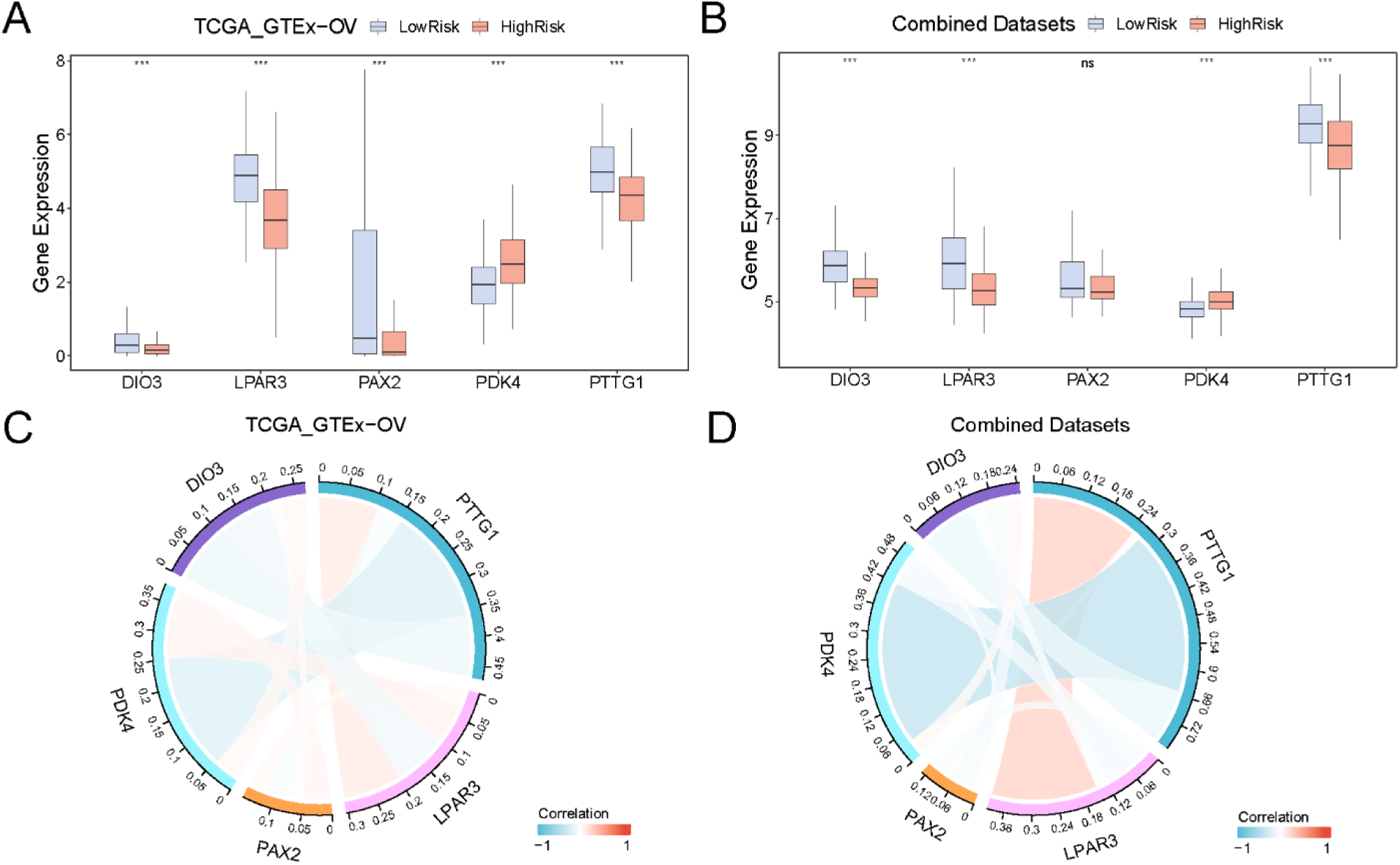
Differential Expression Validation and Correlation Analysis of Model Genes. A-B. Comparative boxplots showing five model genes’ expression difference between high and low-risk groups in TCGA_GTEx-OV (A) and combined datasets (B). C-D. Correlation and chord diagrams of the five model genes in TCGA_GTEx-OV (C) and combined datasets (D). TCGA, The Cancer Genome Atlas; GTEx, Genotype-Tissue Expression; OV, Ovarian cancer; *** represents p < 0.001, highly statistically significant; ns represents p ≥ 0.05, which is not statistically significant. Chord diagrams depicting gene-gene correlations (red: positive; blue: negative; line width scales with |r| value). |r| below 0.3 was weak or no correlation, 0.3-0.5 was a weak correlation, 0.5-0.8 was a moderate correlation, and above 0.8 was a strong correlation.

Cross-Dataset Correlation Patterns Spearman correlation analysis of the five model genes revealed consistent co-expression patterns between the TCGA_GTEx-OV and combined datasets (Fig. 10C-D). In both TCGA_GTEx-OV and combined datasets, *PTTG1* exhibited positive correlation with *LPAR3* and negative correlations with *PDK4* and *DIO3*. *PTTG1* emerged as a central hub in three significant pairs.

### 3.11 Identification of Molecular Subtypes and Correlation Analysis of Model Genes

Consensus clustering analysis based on five model genes (*PTTG1*, *LPAR3*, *PAX2*, *PDK4*, *DIO3*) identified two molecularly distinct subtypes: Subtype 1 (Cluster 1, n = 302) and Subtype 2 (Cluster 2, n = 72) (Fig. 11A-C). The 3D t-SNE visualization (Fig. 11D) showed almost complete spatial separation between the two subtypes, validating clustering robustness. A heatmap of model gene expression (Fig. 11E) revealed that *PAX2* was predominantly downregulated in Subtype 1 and upregulated in Subtype 2, suggesting its role as a subtype-defining marker.

**Fig.11.**
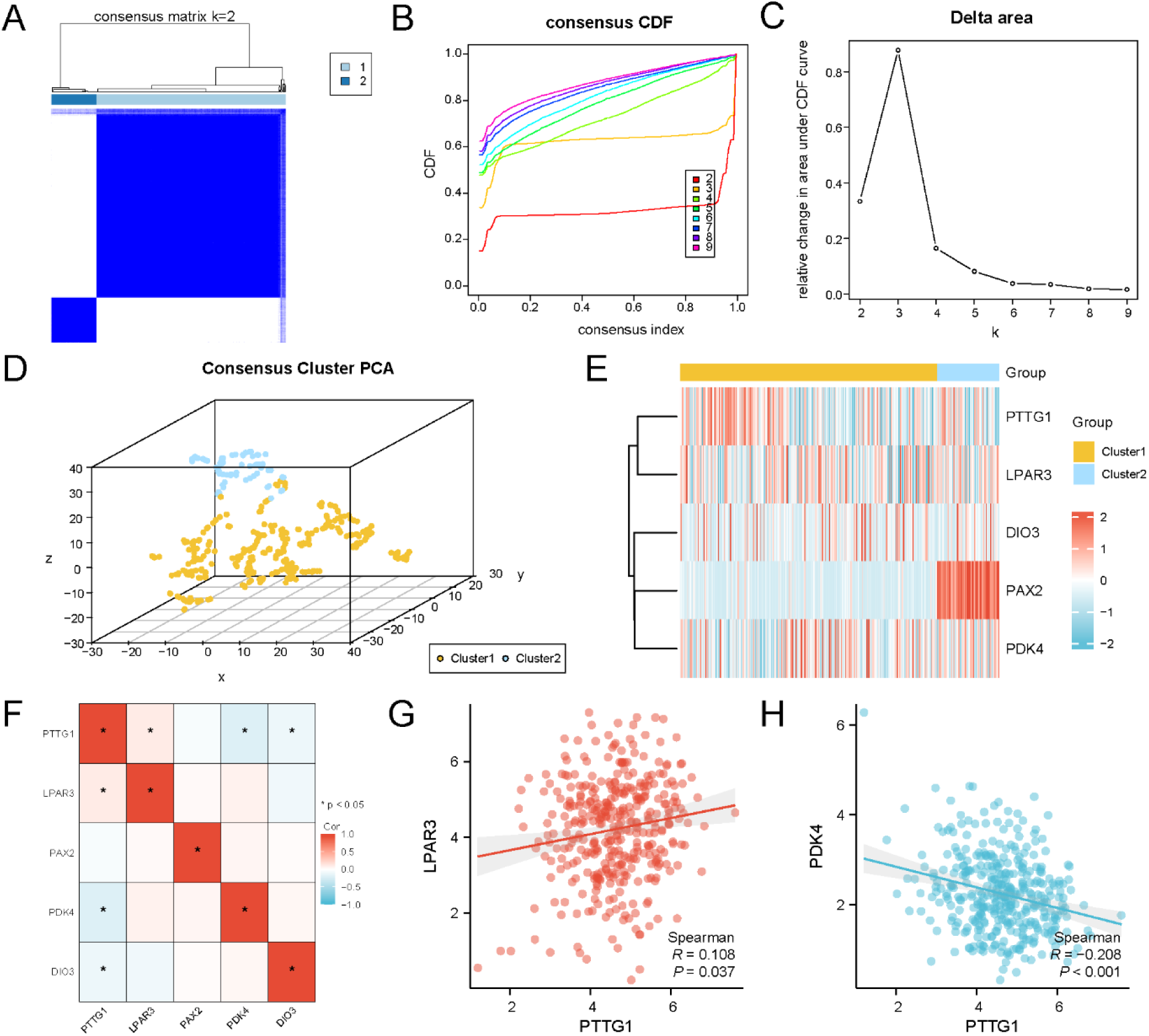
Identification of Ovarian Cancer Subtypes and Correlation Analysis of Model Genes. A. Consensus matrix of k=2 consensus clustering based on five model genes. B-C. Consensus cumulative distribution function (CDF) curve (B) and Delta area plot (C) of consensus clustering analysis. D. 3D t-SNE visualization showing spatial separation of the two subtypes (yellow = Subtype 1, blue = Subtype 2). E. Heatmap of model gene expression across subtypes, with red/blue representing up/downregulation. F. Correlation heatmap of model genes, with red/blue indicating positive/negative correlations; gene pairs marked with * are significantly associated (p < 0.05). G-H. Scatter plots of the top positive (G) and negative (H) correlations, depicting associations between PTTG1-LPAR3 and PTTG1-PDK4, respectively. |r| below 0.3 was weak or no correlation, 0.3-0.5 was a weak correlation, 0.5-0.8 was a moderate correlation, and above 0.8 was a strong correlation.

A correlation heatmap of model genes (Fig. 11F) identified three significantly associated gene pairs (marked with *, p < 0.05): PTTG1-LPAR3, PTTG1-PDK4, and PTTG1-DIO3, consistent with chord diagram results. The top positive (PTTG1-LPAR3, r = 0.108, p = 0.037) and negative (PTTG1-PDK4, Spearman r = −0.208, p < 0.001) correlations were visualized using scatter plots (Fig. 11G-H), providing quantitative evidence for co-regulatory mechanisms.

### 3.12 Immune Infiltration Analysis between High- and Low-Risk Groups

Using the expression matrix from the TCGA_GTEx-OV database, we quantified the infiltration abundance of 28 immune cell types through the ssGSEA algorithm. The comparative analysis revealed significant differences in immune cell infiltration patterns between risk groups. The comparative boxplot (Fig. 12A) demonstrated statistically significant differences (p < 0.05) in seven immune cell types between high- and low-risk OV samples: Activated CD4+ T cells, CD56dim natural killer cells, eosinophils, immature dendritic cells, memory B cells, neutrophils, and type 2 T helper cells.

**Fig.12.**
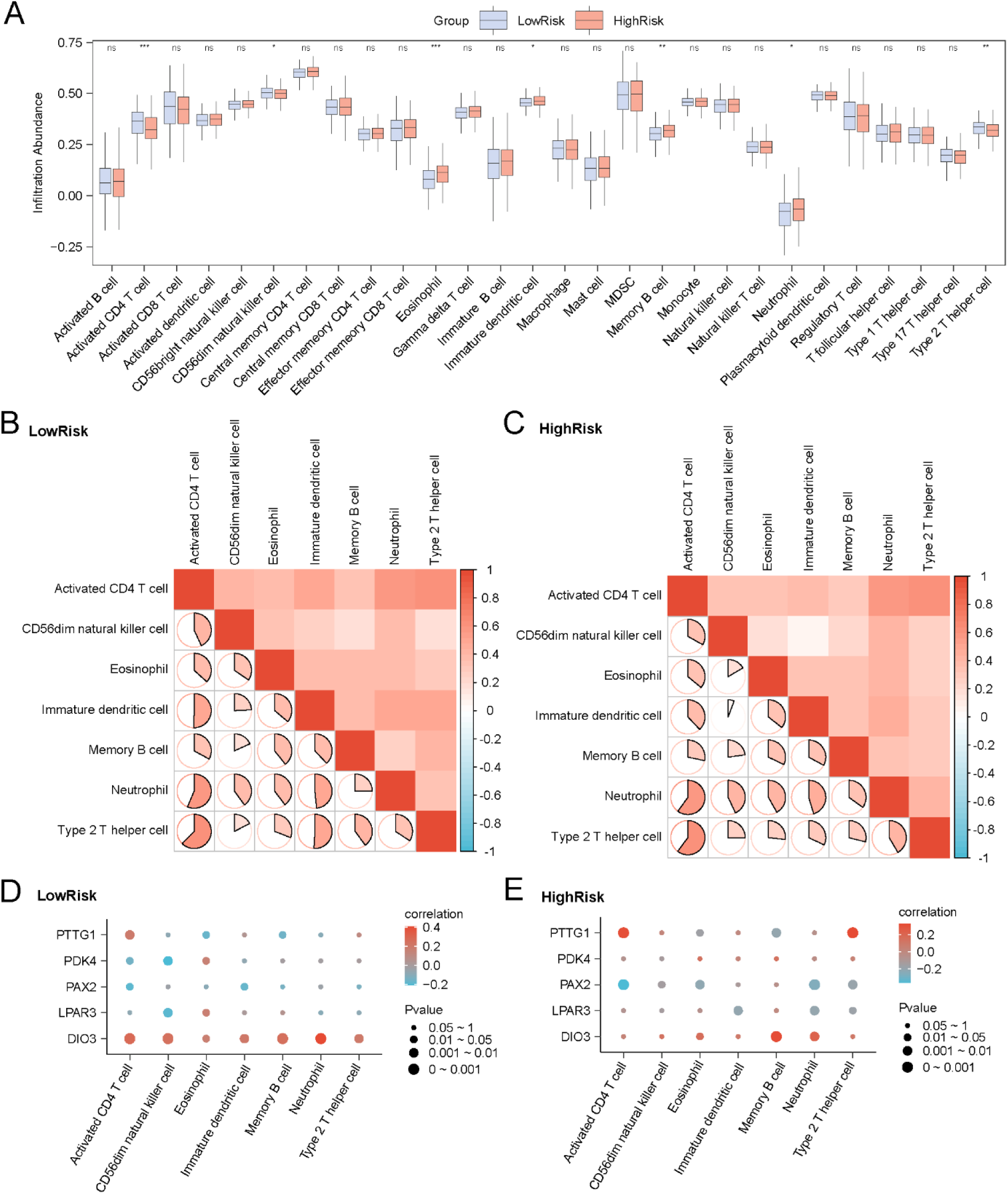
Risk Groups Immune Infiltration Analysis by ssGSEA Algorithm. A. Comparative boxplot of immune cell infiltration between low- and high-risk OV groups in the TCGA_GTEx-OV dataset (blue = low-risk group, red = high-risk group). B-C. Correlation heatmaps of immune cell infiltration abundance in low-risk (B) and high-risk (C) OV groups. D-E. Bubble plots showing correlations between model genes and immune cell infiltration abundance in low-risk (D) and high-risk (E) OV groups. OV, Ovarian cancer; TCGA, The Cancer Genome Atlas; GTEx, Genotype-Tissue Expression; ssGSEA, single-sample Gene Set Enrichment Analysis. ns stands for p ≥ 0.05, not statistically significant; * represents p < 0.05, statistically significant; ** represents p < 0.01, highly statistically significant; *** represents p < 0.001 and highly statistically significant. |r| below 0.3 was weak or no correlation, between 0.3 and 0.5 was a weak correlation, between 0.5 and 0.8 was a moderate correlation, and above 0.8 was a strong correlation. Red is a positive correlation, blue is a negative correlation, and the depth of the color represents the strength of the correlation.

Subsequent correlation analysis (Fig. 12B-C) revealed distinct immune cell interaction patterns between risk groups. In low-risk OV samples, immune cells showed strong positive correlations, with the most significant association observed between type 2 T helper cells and activated CD4+ T cells (r = 0.62, p < 0.05; Fig. 12B). Similarly, high-risk samples exhibited predominantly positive correlations, particularly between activated CD4+ T cells and neutrophils (r = 0.60, p < 0.05; Fig. 12C).

Furthermore, bubble plots illustrating the correlation between model genes and immune cell infiltration (Fig. 12D-E) identified specific gene-immune cell associations. The results showed that the *DIO3* gene and neutrophils had the strongest significant positive correlation in the low-risk samples (r = 0.413, p < 0.05; Fig. 12D), while the *PTTG1* gene and type 2 T helper cells had the strongest significant positive correlation in the high-risk group (r = 0.324, p < 0.05; Fig. 12E).

### 3.13 Immunotherapy Biomarker Assessment: TIDE, MSI, and TMB

Given the emerging importance of immunotherapy in cancer treatment, we assessed immunotherapy sensitivity using the TIDE algorithm and MSI and TMB differences between risk groups in the TCGA_GTEx-OV dataset. Grouped violin plots (Fig. 13) revealed highly statistically significant differences (p < 0.001) in TIDE immunotherapy scores between groups, with low-risk samples showing significantly lower scores than high-risk samples. This suggests potentially better immunotherapy responsiveness in the low-risk group.

**Fig.13.**
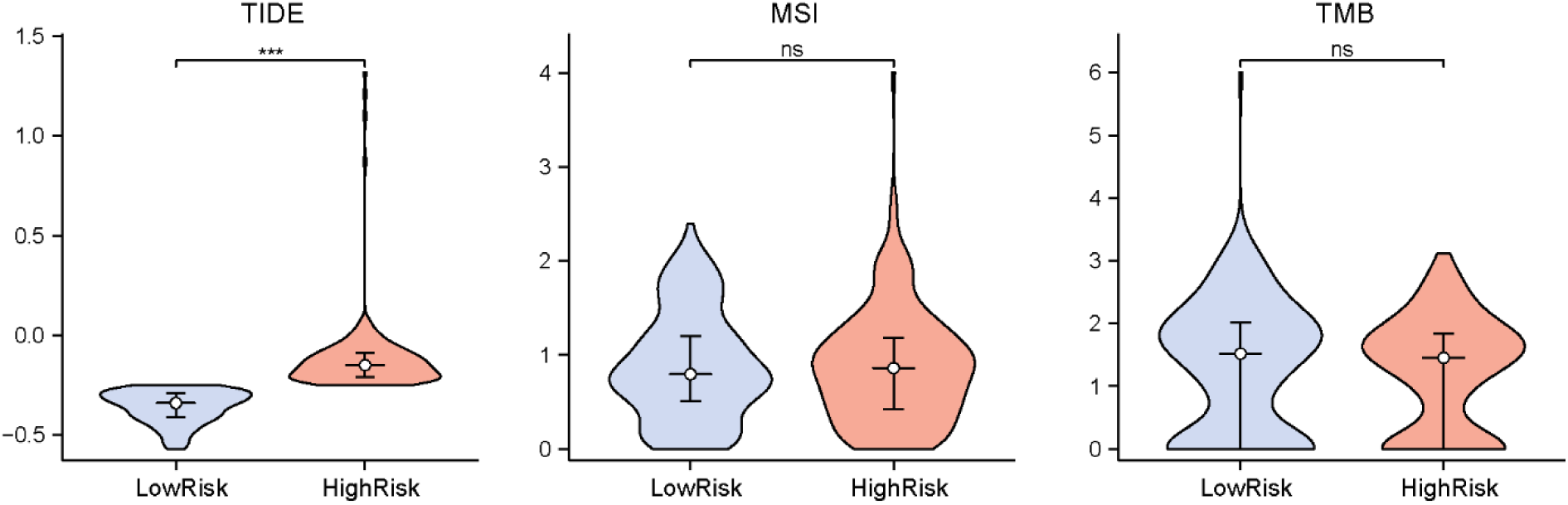
TIDE, MSI, and TMB Analysis. Grouped violin plots of TIDE immunotherapy scores, MSI scores, and TMB scores between high- and low-risk OV groups in the TCGA_GTEx-OV dataset (blue = low-risk group, red = high-risk group). TIDE, Tumor Immune Dysfunction and Exclusion; MSI, Microsatellite Instability; TMB, Tumor Mutation Burden. ns stands for p ≥ 0.05, not statistically significant; *** represents p < 0.001, highly statistically significant.

## 4. Discussion

OV remains the most lethal gynecologic malignancy (1). Due to the lack of reliable early detection tools, most patients are diagnosed at advanced stages (3). Despite the clinical achievements of progressive treatment modalities, high mortality rates persist. The predictive and prognostic value of classic OV biomarkers, such as carbohydrate antigen 125 (CA125) and human epididymis protein 4 (HE4), remains controversial (4), and no reliable tools currently exist for prognostic risk assessment. Metabolic reprogramming is a hallmark of cancer and plays a critical role in tumorigenesis and progression (8). This process not only represents a potential therapeutic target but also serves as a promising biomarker for cancer prognosis. It has demonstrated strong prognostic performance in various cancers, including clear cell renal cell carcinoma (45), pancreatic adenocarcinoma (46), head and neck squamous cell carcinoma (47), lung adenocarcinoma (48), and hepatocellular carcinoma (48). Therefore, leveraging metabolic reprogramming to develop valuable prognostic signatures holds significant clinical importance for OV prognostic risk stratification and personalized treatment strategies. In this study, we conducted bioinformatics analysis to identify MRRDEGs in OV and constructed an effective prognostic risk model based on these MRRDEGs.

The understanding of genomic alterations in OV-associated MRRDEGs remains limited. Our study conducted a systematic analysis of these alterations. Consistent with previous reports, the highest frequency of SM of all genes in OV was *TP53* (94%) (49). The mutation type of SM was mainly SNP, with missense mutations constituting the majority. Among the 64 MRRDEGs, *MUC16* showed the highest mutation rate (8%). *MUC16*, also known as CA125, is expressed in OV epithelial cells and is a well-established biomarker for OV screening (50). Studies have demonstrated that *MUC16* plays a crucial role in cellular adhesion, signaling pathways, and immune regulation, thereby promoting tumor cell proliferation, invasion, and immune evasion (51). CNVs might arise from errors in genome recombination or replication and could influence gene expression through multiple mechanisms (52). CNVs were detected in 52 of the 64 MRRDEGs, which might result in aberrant expression of metabolic genes, consequently leading to abnormal cell growth and promoting tumorigenesis. The high frequency of CNVs in OV suggests that metabolic genes might significantly contribute to disease pathogenesis and could serve as potential therapeutic targets.

Metabolic reprogramming serves as a critical mechanism enabling cancer cells to acquire proliferative advantages, resist apoptosis, and adapt to the harsh TME (14, 53). Our GO and KEGG pathway enrichment analyses revealed that the 64 MRRDEGs were predominantly associated with cell cycle regulation and mitotic biological processes. These findings were consistent with previous reports that tumor cells evade cell cycle regulation and proliferate rapidly through metabolic reprogramming (54). Such dysregulated cell cycle progression might reduce the effective time window for chemotherapeutic agents, potentially contributing to tumor drug resistance (55). Previous studies have reported that cancer cells could bypass cellular senescence and continue to proliferate by altering the expression or function of cell cycle regulatory proteins such as p53 (56). The observed enrichment of MRRDEGs in senescence-related pathways suggests that tumor cells might modulate the expression of these genes during their evasion of senescence and subsequent influence on neighboring normal cells. In addition, the MRRDEGs were also enriched in the p53 signaling pathway. In OV, the *TP53* gene is one of the most frequently mutated genes (57). *TP53* mutations not only impair the normal regulatory functions of the cell cycle but also disrupt the regulation of cellular metabolism (58–60). This dysfunction ultimately results in the uncontrolled proliferation of tumor cells. Metabolic reprogramming of OV cells might also have feedback regulation on the p53 signaling pathway, further affecting the activity and stability of p53 (58).

Based on the MRRDEGs, we developed an effective prognostic risk model including five key genes for OV. These genes not only stratify OV patients into distinct risk groups but also reveal critical biological drivers of disease progression. *PDK4* emerged as the predominant risk factor in our model, consistent with its well-characterized role in promoting glycolytic dependency (Warburg effect) in tumor cells through inhibition of the pyruvate dehydrogenase complex (PDH) (61, 62). Accumulating clinical evidence confirms *PDK4* as a critical metabolic biomarker and oncogene in multiple cancer types (63). In OV specifically, *PDK4* overexpression correlates significantly with metastasis, chemotherapy resistance, and poor prognosis (63, 64). The pro-invasive activity of *PDK4* appears to be mediated through activation of the STAT3/AKT/NF-κB/IL-8 signaling cascade (64). Additionally, emerging research reveals complex regulatory networks involving multiple upstream molecules that modulate *PDK4* expression and consequently influence OV progression (65–67). These findings highlight the role of *PDK4* as a potential therapeutic target and prognostic biomarker in OV.

*PTTG1* has emerged as a critical oncogene in multiple malignancies, with recent studies highlighting its multifaceted role in OV progression (68, 69). Elevated *PTTG1* expression in OV significantly contributes to tumor angiogenesis, M2 macrophage polarization, and epithelial-mesenchymal transition (EMT) (70, 71). Meanwhile, *DIO3*, a thyroid hormone-inactivating enzyme, plays a pivotal role in cellular metabolism regulation (72). The study by Hou et al. (73) revealed that reduced *DIO3* expression correlates with unfavorable OV patient outcomes, including shorter OS and progression-free survival (PFS). Notably, *DIO3* expression patterns showed significant associations with tumor-associated macrophage (TAM) markers and various T-cell subsets in OV. Moskovich et al. (74) further elucidated the context-dependent nature of *DIO3*, suggesting its functional diversity across different OV subtypes, stages, and histological grades. The tumor suppressive potential of the DLK1-DIO3 locus was further elucidated by Cui et al. (75), who identified miR-134-3p as a key regulator that inhibits OV stem cell proliferation and invasion through RAB27A targeting. *LPAR3* contributes to OV progression through multiple mechanisms. Existing evidence confirms its regulation of tumor proliferation via the miR-198/MET axis (76), while emerging data implicate its potential role in cisplatin resistance through miR-634/PDK1 signaling (77).

In our study, *PAX2* effectively classified OV cases into two molecular subtypes, underscoring its significance in tumor subtyping. Research has found that *PAX2* presents a unique expression profile in OV pathogenesis. While absent in normal ovarian surface epithelium, it becomes aberrantly activated in 61% of OV cell lines (78, 79). Experimental evidence supports *PAX2’s* multifaceted role in tumor progression through: (1) angiogenesis promotion, (2) enhanced cancer cell proliferation and invasion, and (3) fatty acid metabolic reprogramming (80, 81). Clinically, elevated *PAX2* expression significantly associates with adverse outcomes in serous OV and shows marked upregulation in platinum-resistant cases (81, 82). However, its occasional tumor-suppressive effects in certain OV subtypes underscore the context-dependent nature of its biological functions (80). The prognostic model genes examined in this study demonstrate significant associations with OV outcomes. While the clinical relevance of certain genes aligns with our findings, the complete prognostic potential of others warrants further investigation to fully elucidate their roles in OV pathogenesis and treatment response.

We observed that immune cells critical for tumor suppression, such as activated CD4+ T cells and CD56dim natural killer cells, were significantly more abundant in the low-risk group. Additionally, the infiltration levels of type 2 T helper cells were also elevated in these patients. Traditionally, the type 2 immune response driven by type 2 T helper cells has been considered immunosuppressive and linked to tumor immune evasion (83). However, emerging evidence suggests that the type 2 immune response might instead enhance anti-tumor immunity, challenging conventional understanding (84). For instance, Feng et al. (85) demonstrated that the type 2 cytokine Fc–IL-4 could directly act on terminally exhausted CD8+ T cells in tumors, boosting their glycolysis, survival, and effector function, thereby synergizing with type 1 immunity to exert anti-tumor effects. In contrast, the immune infiltration in the high-risk group indicated poor antitumor function and possibly even immune escape. Collectively, these findings indicate that patients in the low-risk group exhibit a more robust anti-tumor immune response, contributing to their favorable prognosis. Furthermore, correlation analyses between model genes and immune cell infiltration revealed complex interactions, with *DIO3* showing particularly strong positive correlations with most immune cell types. The finding suggests that these genes might play dual roles in metabolic and immune regulatory processes.

To further assess immunotherapy responsiveness, we applied the TIDE algorithm to evaluate the OV cohort. A lower TIDE score indicates a reduced potential for immune evasion, suggesting a higher likelihood of clinical benefit from immunotherapy. Our analysis revealed significantly lower TIDE scores in the low-risk group compared to the high-risk group, implying superior immunotherapy responsiveness in low-risk patients. Consequently, low-risk patients might be optimal candidates for immune checkpoint inhibitors (e.g., PD-1/PD-L1 blockade), potentially achieving better therapeutic outcomes. These findings could facilitate patient stratification for immunotherapy and enhance treatment efficacy.

Our study has some limitations. First, as a retrospective analysis, our findings and the predictive efficacy of the prognostic risk model require further validation with additional clinical samples. Second, potential batch bias might arise from the heterogeneity of public databases (e.g., variations in sample processing protocols). Although normalization was performed, the generalizability of the model in real-world applications still needs further assessment. Finally, our conclusions are based solely on computational biology approaches; future experimental studies are necessary to clarify the molecular mechanisms of the identified genes and evaluate their clinical utility.

## 5. Conclusions

In conclusion, our study systematically integrated multiple database resources and employed comprehensive bioinformatics approaches to elucidate the role of metabolic reprogramming in OV. These analyses have enhanced our understanding of the relationship between metabolic reprogramming and disease pathogenesis, progression, prognosis, and the tumor immune microenvironment in OV. We developed a clinically promising, metabolism-related prognostic risk model that serves as an independent predictive feature for both prognosis and immunotherapeutic response in OV. The risk score could effectively stratify OV patients into subgroups with significant survival differences, demonstrating potential clinical applicability. Our findings not only reveal potential therapeutic targets but also provide new perspectives for OV patient stratification and treatment strategy development.

